# Muscle-specific DNM2 overexpression improves Charcot-Marie-Tooth disease in vivo and reveals a narrow therapeutic window in skeletal muscle

**DOI:** 10.1101/2025.11.25.690395

**Authors:** Marie Goret, Gwenaelle Piccolo, Jocelyn Laporte

## Abstract

Charcot-Marie-Tooth disease (CMT) caused by dominant loss-of-function mutations in *DNM2*, encoding the GTPase dynamin-2, impairs motor and sensory function. However, the respective contributions of muscle and nerve pathology, and the therapeutic potential of increasing DNM2 expression, remain unresolved. We evaluated tissue-targeted and systemic approaches to increase DNM2 in a mouse model carrying the common K562E-CMT mutation. Muscle-specific DNM2 overexpression from embryogenesis in *Dnm2*^K562E/+^ mice ameliorated desmin and integrin mislocalization, membrane trafficking defects, mitochondrial abnormalities, and fibrosis in skeletal muscle, resulting in improved locomotor performance despite persistent muscle atrophy. Conversely, systemic postnatal AAV delivery of human DNM2 increased DNM2 in muscle but failed to transduce nerves, and paradoxically worsened the muscle pathology, producing centronuclear myopathy-like features. These findings reveal a primary pathogenic impact of *DNM2*-CMT mutation within skeletal muscle, independent of nerve involvement. Collectively, they underscore that precise DNM2 dosage is critical for neuromuscular homeostasis and reveal a narrow therapeutic window for safe and effective therapeutic intervention. This paradox, in which efforts to compensate for a loss-of-function neuropathy risk inducing a gain-of-function myopathy, highlights the need for tightly controlled modulation of DNM2 activity in future therapeutic strategies.

## Introduction

Charcot-Marie-Tooth (CMT) disease, one of the most common inherited peripheral neuropathies (prevalence ∼1:2,500), arises from mutations in over 100 genes^1^. Among these, heterozygous dominant mutations in *DNM2*, which encodes the ubiquitously expressed GTPase dynamin-2 that regulates endocytosis, intracellular trafficking, and cytoskeletal dynamics^2–4^, are associated with both intermediate (CMT-DIB; MIM#606482)^5,6^ and axonal (CMT2M)^7^ forms. Approximately ten different pathogenic *DNM2*-CMT mutations have been identified to date, predominantly clustered within the pleckstrin homology domain, responsible for binding membrane phosphoinositides. These mutations are believed to impair lipid binding, leading to reduced GTPase activity and defective membrane fission^8–11^.

Patients with *DNM2-*CMT mutations exhibit slowed nerve conduction velocities, structural nerve abnormalities, progressive muscle weakness and atrophy, impaired coordination, and skeletal deformities^12^. Additional signs may include ptosis, ophthalmoparesis, cataracts, or neutropenia^13–15^. Despite its clinical and genetic heterogeneity, no disease-modifying therapies are approved for CMT^16,17^, highlighting the urgent need for mechanistic studies and targeted interventions.

The *K562E* heterozygous missense mutation is the most common, and the *Dnm2^K^*^562^*^E/+^* mouse model recapitulates motor deficits, muscle atrophy, along with mild peripheral nerve defects^10,18,19^. While the initial characterization of this model suggested a primary myopathy^18^, early nerve involvement, such as reduced g-ratio in sciatic nerves at 8 weeks of age, were also described^10,19^. These findings raise the unresolved question of whether muscle pathology arises secondarily to nerve dysfunction or whether the mutation exert a direct, cell-autonomous effect in skeletal muscle^20^.

Furthermore, while loss-of-function mutations cause CMT, gain-of-function mutations lead to centronuclear myopathy (CNM)^6,21^, a primary muscle disorder characterized by muscle weakness and atrophy, ptosis, ophthalmoplegia, and centralized organelles^6,8,21^. The gain-of-function effect of CNM mutations was confirmed by studies showing that DNM2 overexpression in wild-type (WT) mice induces a CNM-like phenotype^22,23^. Genetic studies in mice combining CMT and CNM mutations have shown that rebalancing DNM2 activity toward normal levels can prevent the development of both diseases^10^. Moreover, reducing DNM2 protein levels in CNM mouse models with different methodologies ameliorated disease phenotypes^24–27^. Conversely, whether increasing DNM2 expression can benefit CMT without inducing CNM-like pathology has never been tested.

To elucidate the muscle contribution to the pathogenesis underlying *DNM2*-CMT and to evaluate the therapeutic potential of DNM2 overexpression, we tested two approaches in the *Dnm2^K^*^562^*^E/+^* CMT mouse model: (1) muscle-specific DNM2 overexpression from embryogenesis, and (2) postnatal delivery of human DNM2 via adeno-associated virus (AAV) vectors. Muscle-specific DNM2 overexpression initiated during embryogenesis improved several behavioral outcomes and most molecular and histological features in the *Dnm2^K^*^562^*^E/+^* model. In contrast, systemic delivery of human DNM2 at birth increased DNM2 levels in muscle but not in nerve tissue, and failed to improve, and in some cases worsened, muscle phenotypes. These findings identify skeletal muscle as a direct contributor to *DNM2*-CMT pathogenesis and demonstrate that both the level and timing of DNM2 expression critically determine therapeutic outcomes. This work provides crucial guidance for the design of safe and effective gene-based interventions and underscores the potential risks of overexpression-based therapeutic approaches.

## Results

### Muscle-specific DNM2 overexpression from embryogenesis improves *Dnm2*-CMT locomotor performance

The heterozygous K562E mutation, which causes CMT in humans, behaves as a loss-of-function allele with an additional dominant-negative effect in mice, as heterozygous *Dnm2* knockout animals do not recapitulate the neuromuscular phenotypes seen in *Dnm2^K^*^562^*^E/+^* mice^18,28^. To investigate the therapeutic potential of increasing WT DNM2 levels, we first intended to cross *Dnm2^K^*^562^*^E/+^* mice with a transgenic line ubiquitously (Ub) overexpressing murine DNM2 (TgDNM2^Ub^) in which DNM2 expression is triggered by expression of the Cre recombinase under the control of the ACTB promoter. However, TgDNM2^Ub^ mice exhibited perinatal lethality. Although embryos were detected at embryonic day 18.5 (E18.5), no viable pups with this genotype were identified at postnatal day 10 (P10) (Supplementary Fig. 1A-B).

Therefore, as a main affected tissue in the *Dnm2^K^*^562^*^E/+^* mouse is muscle, and to further elucidate the impact of the K562E CMT mutation within this tissue, we restricted the overexpression of DNM2 specifically in striated muscles (SM) upon expression of the Cre recombinase under the control of the muscle-specific MCK promoter, by crossing *Dnm2^K^*^562^*^E/+^* mice with TgDNM2^SM^ transgenic mice. The resulting *Dnm2^K^*^562^*^E/+^*;TgDNM2^SM^ mice were viable, and we assessed their behavioral and locomotor capacities at 8 weeks (Fig. 1A). RT-qPCR showed a 3.2- and 2.1-fold increase in *Dnm2* RNA in tibialis anterior (TA) muscle from *Dnm2^+/+^*;TgDNM2^SM^ and *Dnm2^K^*^562^*^E/+^*;TgDNM2^SM^ mice respectively, compared to the endogenous DNM2 levels in non-overexpressing controls, while western blotting showed a 3.8- and 4.8-fold increase in protein expression (Supplementary Fig. 2A-B). Some inter-animal variability was observed, likely reflecting mosaic MCK-Cre activity and variable recombination efficiency.

**Figure 1.**
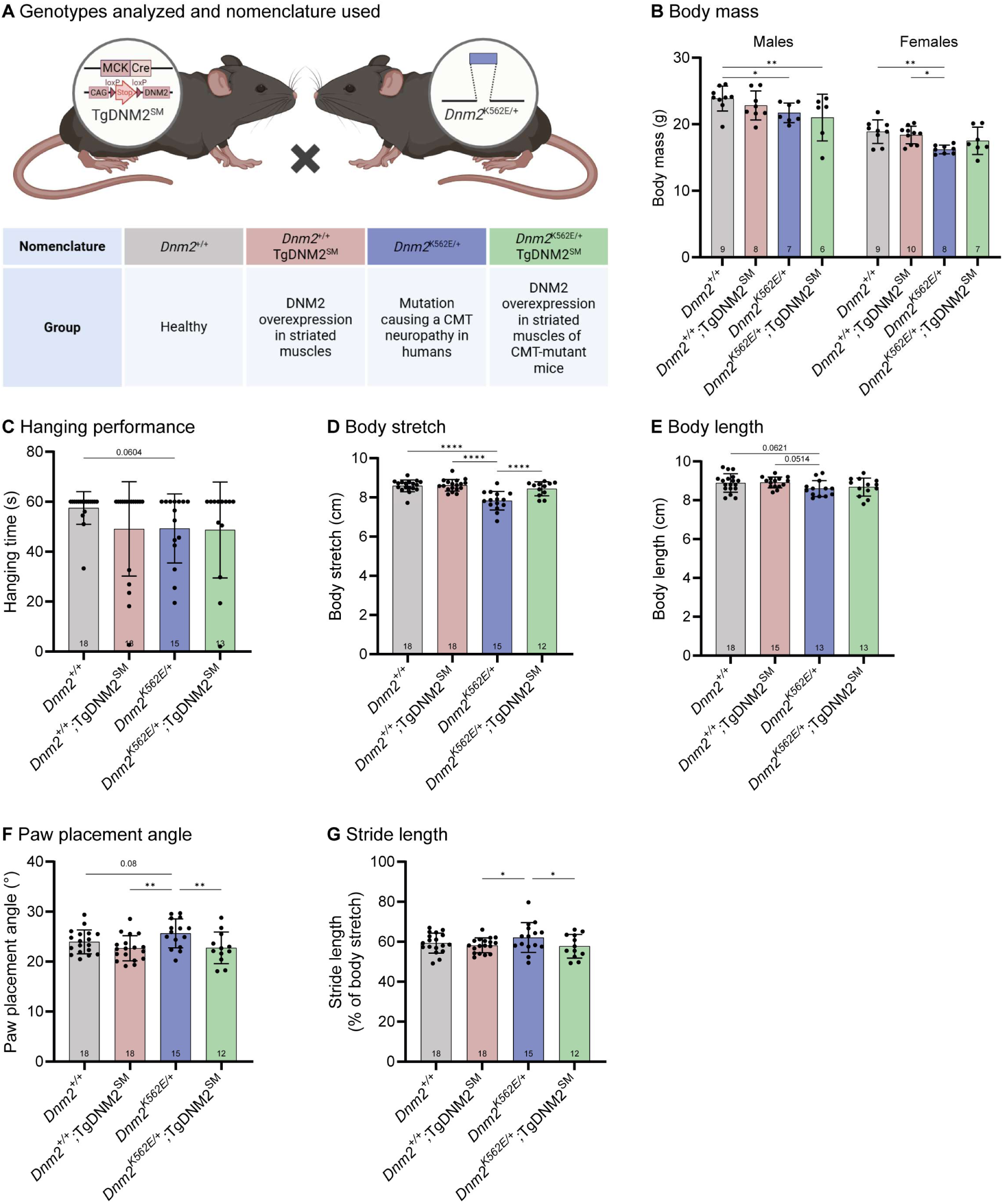
Muscle-specific DNM2 overexpression from embryogenesis improves *Dnm2*-CMT locomotor performance. **(A)** Genotypes analyzed and nomenclature used. Created with BioRender. **(B)** Body mass of males (6≤n≤9) and females at 8w (7≤n≤10). **(C)** Hanging test performance at 8w. Maximum hanging time= 60s (13≤n≤18). **(D)** Body stretch (nose to tail base length) measured during treadmill walking at 8w (12≤n≤18). **(E)** Body length (nose to tail base) measured after euthanasia at 8w (13≤n≤18). **(F)** Angle of feet between paw and body line during treadmill walking (12≤n≤18). **(G)** Stride (=length of a step) during treadmill walking normalized to body stretch (12≤n≤18). Each dot represents a mouse. Values are represented as mean ± SD, *p<0.05, **p<0.01, ****p<0.0001. (B, D-G) ANOVA test. (C) Kruskal-Wallis test.

*Dnm2^K^*^562^*^E/+^*mice showed reduced body mass compared to controls, which was not rescued in males and showed a tendency toward improvement in females following DNM2 overexpression (not different from disease nor control groups) (Fig. 1B). As no sex differences were reported in the *Dnm2*^K562E/+^ mouse model^10^, data from males and females were pooled for all analyses, except for body mass. Although not statistically significant, the *Dnm2^K^*^562^*^E/+^*mice exhibited a decrease in hanging time (p = 0.0604, Fig. 1C), which was not modified upon DNM2 overexpression. When walking on the treadmill, *Dnm2^K^*^562^*^E/+^* mice appeared smaller, as indicated by a significant reduction in body stretch (Fig. 1D), likely attributable to smaller overall body size (Fig. 1E, not significant). Of note, body stretch was fully rescued following DNM2 overexpression, despite no increase in body length, suggesting a functional improvement in muscle tone rather than overall growth. Increased paw placement angle and stride length in *Dnm2^K^*^562^*^E/+^*mice pointed to coordination impairments, and were rescued by muscle-specific DNM2 overexpression (Fig. 1F-G).

### Muscle-specific DNM2 overexpression does not improve *Dnm2*-CMT muscle atrophy

To better understand the motor improvement observed, we analyzed muscle mass and histology. Both TA and Soleus muscle masses were decreased in *Dnm2^K^*^562^*^E/+^*mice compared to controls (Supplementary Fig. 2C-D). DNM2 overexpression did not improve TA mass and showed only a non-significant trend toward improvement in Soleus mass. Histological analysis of TA muscle revealed myofiber hypotrophy, with a 1.2-fold higher proportion of small fibers compared to controls, which was not rescued by DNM2 overexpression (Supplementary Fig. 2E-G). No significant increase in centralized nuclei was observed following DNM2 overexpression (Supplementary Fig. 2H).

Overall, muscle-specific DNM2 overexpression improved or fully rescued several locomotor defects including the body stretch, paw placement and stride length of *Dnm2^K^*^562^*^E/+^* mice but did not rescue muscle atrophy.

### Muscle-specific DNM2 overexpression partially improves *Dnm2*-CMT muscle organization

Since DNM2 overexpression improved locomotor function but not muscle atrophy in *Dnm2^K^*^562^*^E/+^* mice, we next examined the molecular and cellular bases of this apparent discrepancy. Desmin, a key cytoskeletal protein that was shown to interact^29^ and colocalize with DNM2^23^, and to be disrupted by *DNM2* mutations or depletion^10,26,29^, was examined to assess muscle organization. Desmin immunofluorescence in transversal TA sections revealed altered intensity distributions (Fig. 2A, upper panel, white arrows). Control muscles had a high proportion of low-intensity fibers. *Dnm2^K^*^562^*^E/+^* muscles had ∼20% of fibers exhibiting high intensity (mean gray value >100), compared to ∼7% in WT controls, though this difference was not statistically significant (Fig. 2B-C), suggesting impaired protein turnover. This higher proportion of fibers with high desmin intensity was not rescued in *Dnm2^K^*^562^*^E/+^*;TgDNM2^SM^, while *Dnm2^+/+^*;TgDNM2^SM^ muscles showed a shift toward higher intensities. The tendency of desmin mislocalization in *Dnm2^K^*^562^*^E/+^* muscle correlated with increased desmin protein levels, which were completely normalized following DNM2 overexpression (Fig. 2D, Supplementary Fig. 3A).

**Figure 2.**
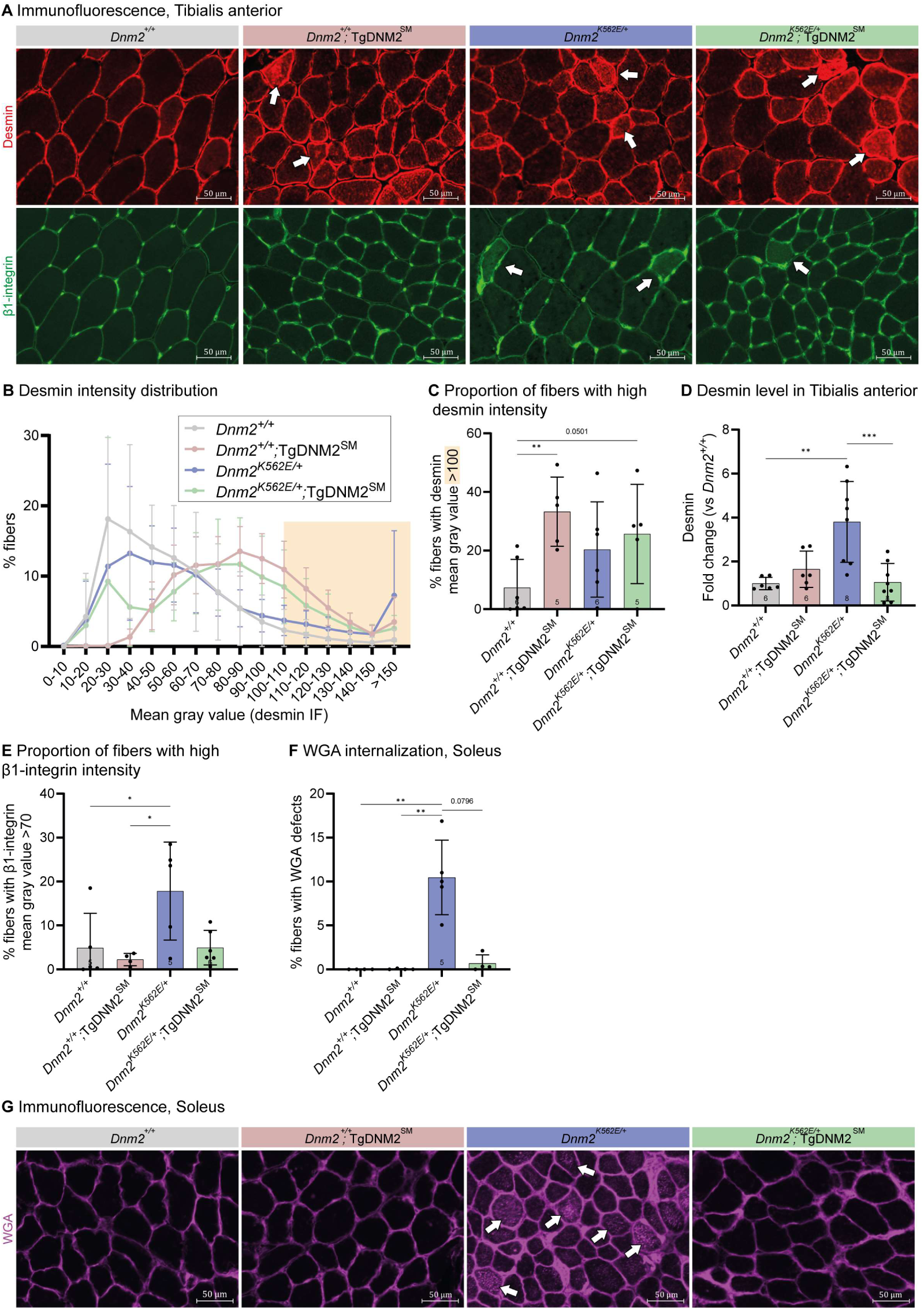
Muscle-specific DNM2 overexpression partially improves *Dnm2*-CMT muscle organization. **(A)** Immunolabeling of desmin (top panel) and β1-integrin (second panel) in transversal Tibialis anterior (TA) sections. Arrows indicate examples of fibers presenting an abnormal central accumulation of the staining. Scale bar= 50 µm. **(B)** TA fibers distribution based on their desmin fluorescence intensity (higher gray value, brighter the fiber) (5≤n≤6). **(C)** Proportion of fibers with high desmin intensity (mean gray value >100) (5≤n≤6). **(D)** Quantification of Desmin protein level in TA at 8w, normalized to Ponceau S staining, depicted in supplementary material (6≤n≤8). **(E)** Proportion of fibers with high integrin intensity (mean gray value >70) (4≤n≤6). **(F)** Proportion of fibers presenting an abnormal localization of WGA in Soleus sections at 8w (4≤n≤5). Each dot represents a mouse. Values are represented as mean ± SD, *p<0.05, **p<0.01, ***p<0.001. (D) ANOVA test. (C, E-F) Kruskal-Wallis test.

Similarly, β1-integrin, an adhesion molecule that traffics in a dynamin-dependent pathway and links the cytoskeleton to the extracellular matrix (ECM)^30,31^, showed abnormal intracellular staining in some *Dnm2^K^*^562^*^E/+^*fibers (Fig. 2A, lower panel, white arrows). Analysis of fibers with high β1-integrin staining intensity (mean gray value >70) revealed an increase in *Dnm2^K^*^562^*^E/+^*mice (18%) compared to controls (5%) (Fig. 2E). Overexpression of DNM2 in *Dnm2^K^*^562^*^E/+^*;TgDNM2^SM^ muscles restored the proportion of high-intensity fibers to control levels (5%). In addition, immunofluorescence of wheat-germ-agglutinin (WGA), which labels surface glycoproteins and glycolipids, revealed abnormal internal staining inside ∼10% of fibers in *Dnm2^K^*^562^*^E/+^* sections in Soleus muscle (Fig. 2F-G, white arrows). WGA fluorescence has been previously used to assess membrane internalization in muscle cells^32^. Together with integrin findings, these results support impaired membrane trafficking in *Dnm2^K^*^562^*^E/+^*muscle. WGA internalization was fully rescued in *Dnm2^K^*^562^*^E/+^*;TgDNM2^SM^ overexpressing DNM2.

Prior transcriptomic and histological analyses of Soleus muscle revealed increased ECM deposition in *Dnm2^K^*^562^*^E/+^* mice^18,19^, a finding also observed in *DNM2*-CMT patient^14^. We thus assessed collagen VI expression and found increased collagen thickness between TA muscle fibers in *Dnm2^K^*^562^*^E/+^*mice (Supplementary Fig. 3B, D), confirmed by Masson’s Trichrome staining showing fibrosis in the Soleus muscle (Supplementary Fig. 3C, E, white arrows). While DNM2 overexpression did not improve fibrosis in Soleus, it fully normalized collagen thickness in TA muscle.

Overall, muscle-specific DNM2 overexpression corrected key defects in cytoskeletal organization, membrane dynamics, and extracellular matrix structure in *Dnm2^K^*^562^*^E/+^* mice, that potentially explain the rescue of body stretch, paw placement and stride length despite a conserved muscle atrophy.

### Muscle-specific DNM2 overexpression fully rescues *Dnm2-*CMT mitochondrial dysfunction

Previous transcriptomic and molecular analyses of Soleus muscle revealed mitochondrial dysregulation in *Dnm2^K^*^562^*^E/+^* mice^18,19^, which may underlie part of the locomotor defects of *DNM2*-CMT. We thus assessed the expression of electron transport chain complexes by western blot and found that Complexes I, II, and III were downregulated in *Dnm2^K^*^562^*^E/+^* mice (Fig. 3A-B). DNM2 overexpression fully restored the levels of the affected complexes. Complex I (NADH dehydrogenase) reduction in *Dnm2^K^*^562^*^E/+^* mice correlated with a decreased NADH staining intensity in muscle fibers and presence of abnormal internal staining in some fibers (Fig. 3C-E). DNM2 overexpression normalized NADH staining intensity and improved staining localization. Complex II (succinate dehydrogenase) reduction did not associate with SDH staining abnormalities, as previously reported in TA^19^. In contrast, Complex III (ubiquinol-cytochrome c oxidoreductase) reduction was associated with a marked decrease in cytochrome c protein levels (Fig. 3F), which was fully normalized by DNM2 overexpression. Importantly, these changes were not due to reduced mitochondrial content (Fig. 3G). Restored mitochondrial function likely improves muscle energy metabolism, which may contribute to the locomotor improvements observed despite persistent atrophy.

**Figure 3.**
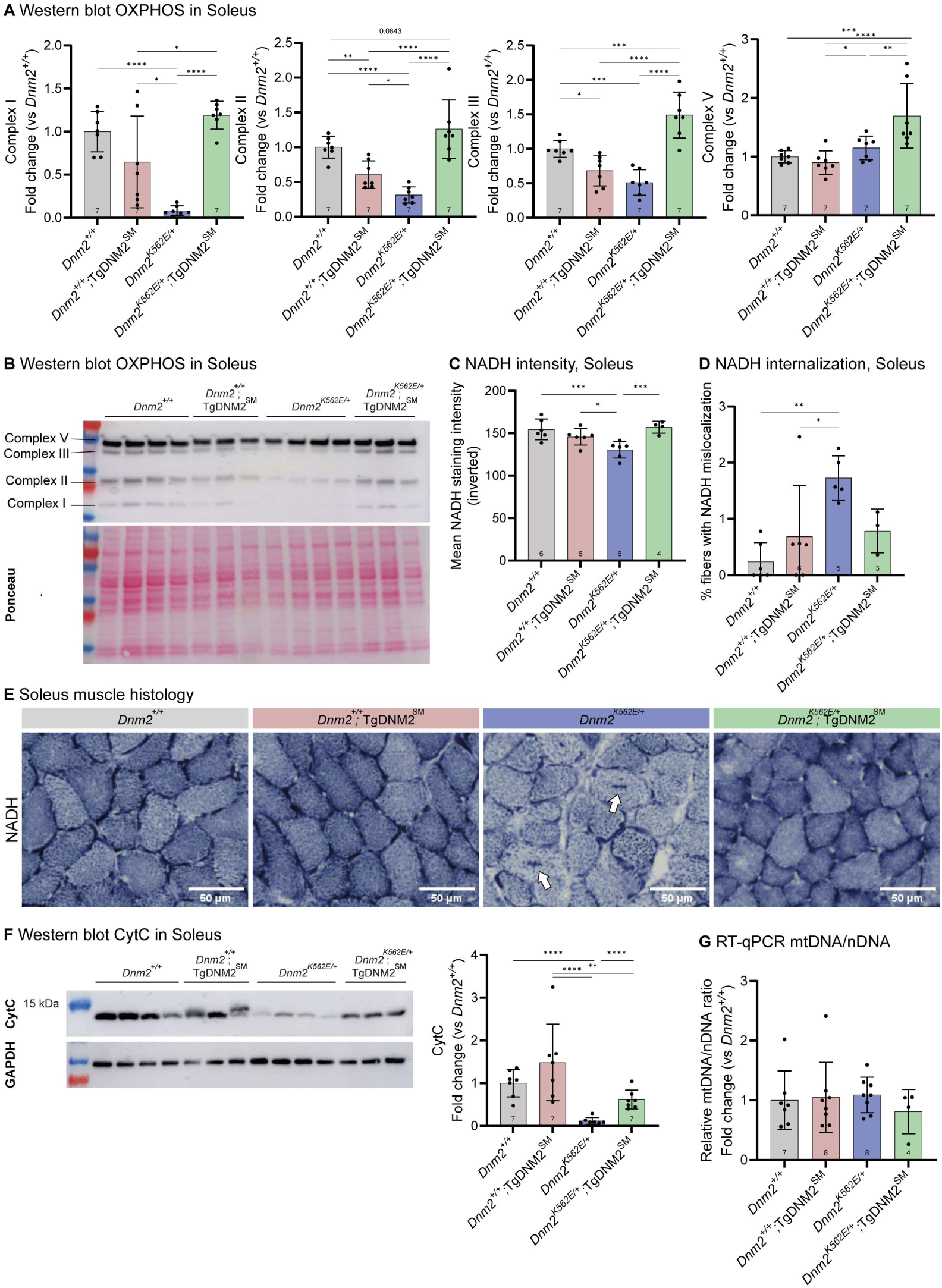
Muscle-specific DNM2 overexpression fully rescues *Dnm2*-CMT mitochondrial dysfunction. **(A)** Quantification of complexes I, II, III, V of the electron transport chain, assessed using the OXPHOS antibody cocktail in Soleus (n=7), and **(B)** representative western blot. **(C)** Mean NADH staining intensity in Soleus muscle fibers, shown as inverted grayscale values (255-raw intensity), so that higher values correspond to darker staining (4≤n≤6). **(D)** Proportion of fibers displaying internalized NADH staining (3≤n≤6). **(E)** Soleus transversal sections stained for nicotinamide adenine dinucleotide (reduced form) (NADH); arrows indicate fibers with abnormal central accumulation of staining. Scale bar = 50 µm. **(F)** Quantification of cytC protein level in Soleus, normalized to Ponceau S staining, depicted in supplementary material (n=7). **(G)** Mitochondrial DNA (mtDNA) to nuclear DNA (nDNA) ratio in Soleus muscle, quantified by RT-qPCR using *Nd1* and *Rsp11* respectively (4≤n≤8). Each dot represents a mouse. Values are represented as mean ± SD, *p<0.05, **p<0.01, ***p<0.001, ****p<0.0001. (A, C, F) ANOVA test. (D, G) Kruskal-Wallis test.

We observed a shift toward faster fiber types in the Soleus in *Dnm2^K^*^562^*^E/+^* mice, with a reduced proportion of type I fibers and an increased proportion of type IIa fibers (Supplementary Fig. 3F). DNM2 upregulation fully rescued the proportion of type I fibers and partially restored type IIa levels. These changes may explain the overall mitochondrial abnormalities observed in *Dnm2^K^*^562^*^E/+^* mice and their rescue following DNM2 upregulation.

In conclusion, muscle-specific DNM2 overexpression (4.8-fold increase vs WT controls, 4.1-fold vs *Dnm2^K^*^562^*^E/+^*) in *Dnm2^K^*^562^*^E/+^*;TgDNM2^SM^ mice led to locomotor improvements but failed to rescue TA muscle atrophy. However, it significantly corrected desmin protein levels and improved membrane trafficking defects (abnormally high integrin intensity and WGA internalization). It also normalized collagen VI thickness in TA and mitochondrial dysfunction in the Soleus.

Overall, these data reveal a primary involvement of skeletal muscle in the *DNM2-*CMT pathology and support early DNM2 augmentation as a potential therapeutic strategy.

### Postnatal DNM2 overexpression does not improve *Dnm2*-CMT whole-body motor performance

As increasing DNM2 level in striated muscles from embryogenesis showed beneficial effects in *Dnm2^K^*^562^*^E/+^*model, we next explored a translational gene therapy approach aimed at preventing disease onset. To reach systemic transduction and assess therapeutic efficacy once embryogenesis is finalized, we administered AAV9 vectors carrying human DNM2 under the ubiquitous CAG promoter via intraperitoneal injection at birth at a dose of 1.0 × 10¹¹ genome copies (gc) per pup and evaluated outcomes at 8 weeks of age (Fig. 4A).

**Figure 4.**
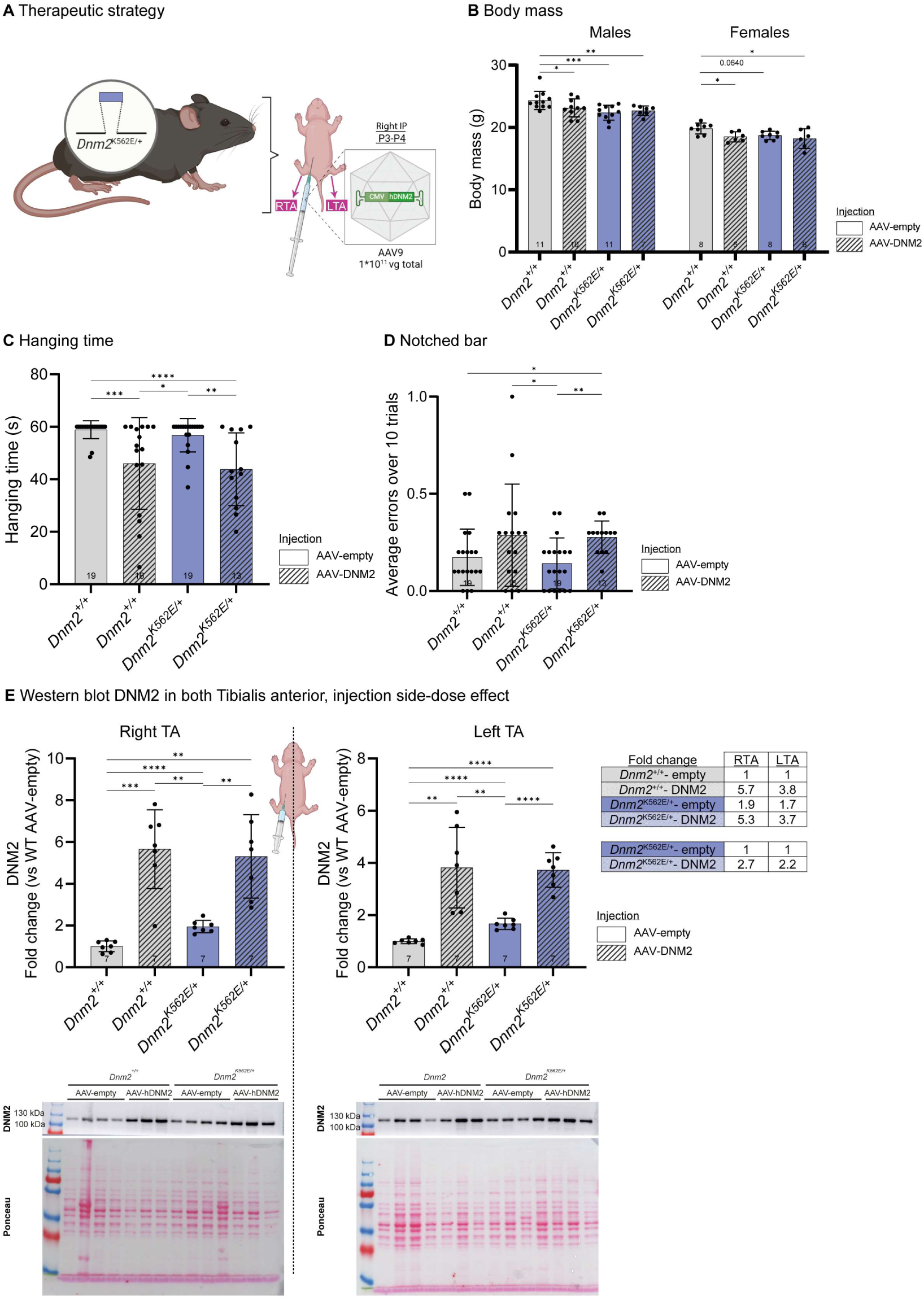
Postnatal DNM2 overexpression does not improve *Dnm2*-CMT whole-body motor performance. **(A)** Experimental set-up. Created with BioRender. **(B)** Body mass of males (7≤n≤11) and females at 8w (6≤n≤8). **(C)** Hanging test performance at 8w. Maximum hanging time= 60s (13≤n≤19). **(D)** Average number of errors (hindlimb slips) when crossing the notched bar over 10 trials (13≤n≤19). **(E)** Representative western blot and quantification of DNM2 protein in right TA (IP-injected side) and left TA (contralateral side), normalized to Ponceau S staining (n=7). The summary table reports the fold change in DNM2 levels for each bodyside, relative to untreated *Dnm2^+/+^* and *Dnm2^K^*^562^*^E/+^* groups. Each dot represents a mouse. Values are represented as mean ± SD, *p<0.05, **p<0.01, ***p<0.001, ****p<0.0001. (B, E) ANOVA test. (C-D) Kruskal-Wallis test.

The 3.6-fold increase of *DNM2* RNA level in *Dnm2^K^*^562^*^E/+^* mice injected with AAV-DNM2 (Supplementary Fig. 4A) did not improve the body mass decrease (Fig. 4B). Postnatal DNM2 overexpression worsened the hanging abilities of *Dnm2^+/+^* and *Dnm2^K^*^562^*^E/+^* mice compared to the untreated groups (Fig. 4C). DNM2 overexpression also induced coordination defects in *Dnm2^K^*^562^*^E/+^* mice, with an increase number of errors when crossing the notched bar (Fig. 4D).

Noteworthy, the intraperitoneal injection was consistently made on the right side in all mice. Western blot analysis showed an average 5.5-fold increase in DNM2 protein levels in the right TA (RTA, IP injection side) of *Dnm2^+/+^* and *Dnm2^K^*^562^*^E/+^* mice, compared to untreated *Dnm2^+/+^* (Fig. 4E). A lower level of overexpression was observed in the contralateral left TA, averaging a 3.8-fold increase. This side-dependent difference in DNM2 expression, resulting from the unilateral intraperitoneal injection, enabled evaluation of two distinct doses within the same animal in subsequent analyses.

### Postnatal DNM2 overexpression does not rescue *Dnm2*-CMT myofiber hypotrophy and promotes CNM histopathology

Postnatal DNM2 delivery worsened motor performance in *Dnm2^K^*^562^*^E/+^* mice. As intraperitoneal injection at birth produced asymmetric expression, we explored whether higher local DNM2 expression (injected right side ∼5.5-fold) could induce myopathic features, consistent with CNM gain-of-function phenotypes, while lower expression (contralateral left side ∼3.8-fold) might be beneficial. We therefore analyzed muscle structure and organization in both hindlimbs.

TA muscle mass, which was reduced in *Dnm2^K^*^562^*^E/+^* mice, was restored to *Dnm2^+/+^* levels on both sides following DNM2 delivery (Supplementary Fig. 4A-B). However, this recovery did not translate into improved muscle fiber morphology. Myofiber hypotrophy persisted, with an elevated proportion of small-diameter fibers in *Dnm2^K^*^562^*^E/+^*mice on both the high-expression (right, ∼5.5-fold) and lower-expression (left, ∼3.8-fold) hindlimbs (Fig. 5A-B), suggesting that the muscle mass increase result from ECM expansion or fat infiltration rather than myofiber hypertrophy.

**Figure 5.**
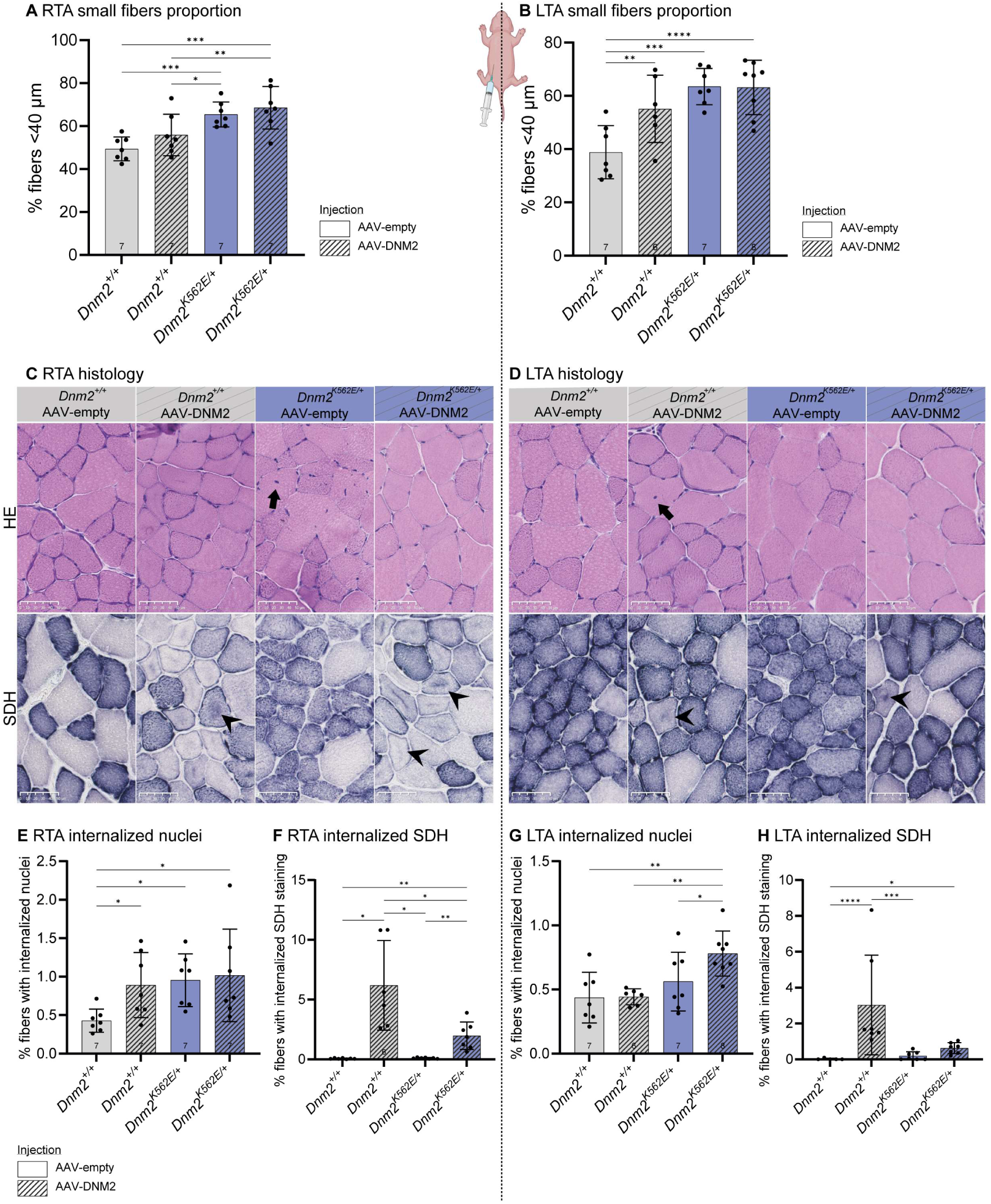
Postnatal DNM2 overexpression does not rescue *Dnm2*-CMT myofiber hypotrophy and promotes CNM histopathology. **(A-B)** Proportion of small fibers (MinFeret<40µm) in **(A)** right TA (RTA, IP-injected side) (n=7) and **(B)** left TA (LTA, contralateral side) (6≤n≤8). **(C–D)** Transversal sections of TA stained with hematoxylin and eosin (HE, top panel), and succinate dehydrogenase (SDH, second panel) in **(C)** right TA (IP-injected side) and **(D)** left TA (contralateral side). Scale bar = 50 µm. Black arrows indicate internalized nuclei; black arrowheads internalized oxidative staining. **(E-F)** Proportion of fibers with: **(E)** internalized nuclei, and **(F)** internalized oxidative staining in right TA (IP-injected side) (6≤n≤7). **(G-H)** Proportion of fibers with: **(G)** internalized nuclei, and **(H)** internalized oxidative staining in left TA (contralateral side) (6≤n≤8). Each dot represents a mouse. Values are represented as mean ± SD, *p<0.05, **p<0.01, ***p<0.001, ****p<0.0001. (A-G) ANOVA test. (H) Kruskal-Wallis test.

Histological analysis revealed abnormalities linked to DNM2 overexpression. In particular, hematoxylin-eosin (HE) staining revealed that nuclear mispositioning (Fig. 5C-D, first panel, black arrows) was exacerbated in *Dnm2^+/+^* mouse TA receiving the high dose (Fig. 5E) and in *Dnm2^K^*^562^*^E/+^* mouse TA receiving the lower dose (Fig. 5G). Similarly, succinate dehydrogenase (SDH) staining revealed disorganized mitochondrial positioning (Fig. 5C-D, second panel, black arrowheads) induced by DNM2 overexpression in both genotypes, particularly at higher doses (Fig. 5F, H). The myofiber hypotrophy combined with the increase in nuclear and mitochondria misposition recapitulated the main CNM histopathological hallmarks following postnatal DNM2 overexpression.

### Postnatal DNM2 overexpression fails to restore desmin and integrin localization in *Dnm2-*CMT muscle

We next assessed desmin and β1-integrin localization by immunofluorescence, given their previously observed mislocalization in *Dnm2^K^*^562^*^E/+^* muscle and rescue following embryonic DNM2 overexpression. Desmin accumulated abnormally in ∼31% of *Dnm2^K^*^562^*^E/+^* TA fibers, compared to ∼11% in controls, and this defect was not rescued by either dose of AAV-DNM2 (Fig. 6A-B, first panel, red arrows; Fig. 6C-D). Similarly, integrin was mislocalized inside myofibers in ∼18% of *Dnm2^K^*^562^*^E/+^* fibers versus ∼4% in *Dnm2^+/+^*. AAV-DNM2 worsened this defect, the high dose increased mislocalization in both genotypes as the low dose in *Dnm2^+/+^*, without benefit in *Dnm2^K^*^562^*^E/+^* (Fig. 6A-B, second panel, red arrowheads; Fig. 6E-F). As postnatal intraperitoneal AAV-DNM2 injection did not increase DNM2 levels in the sciatic nerve at 8 weeks on either body side, nerve analysis was not pursued (Supplementary Fig. 4D-E).

**Figure 6.**
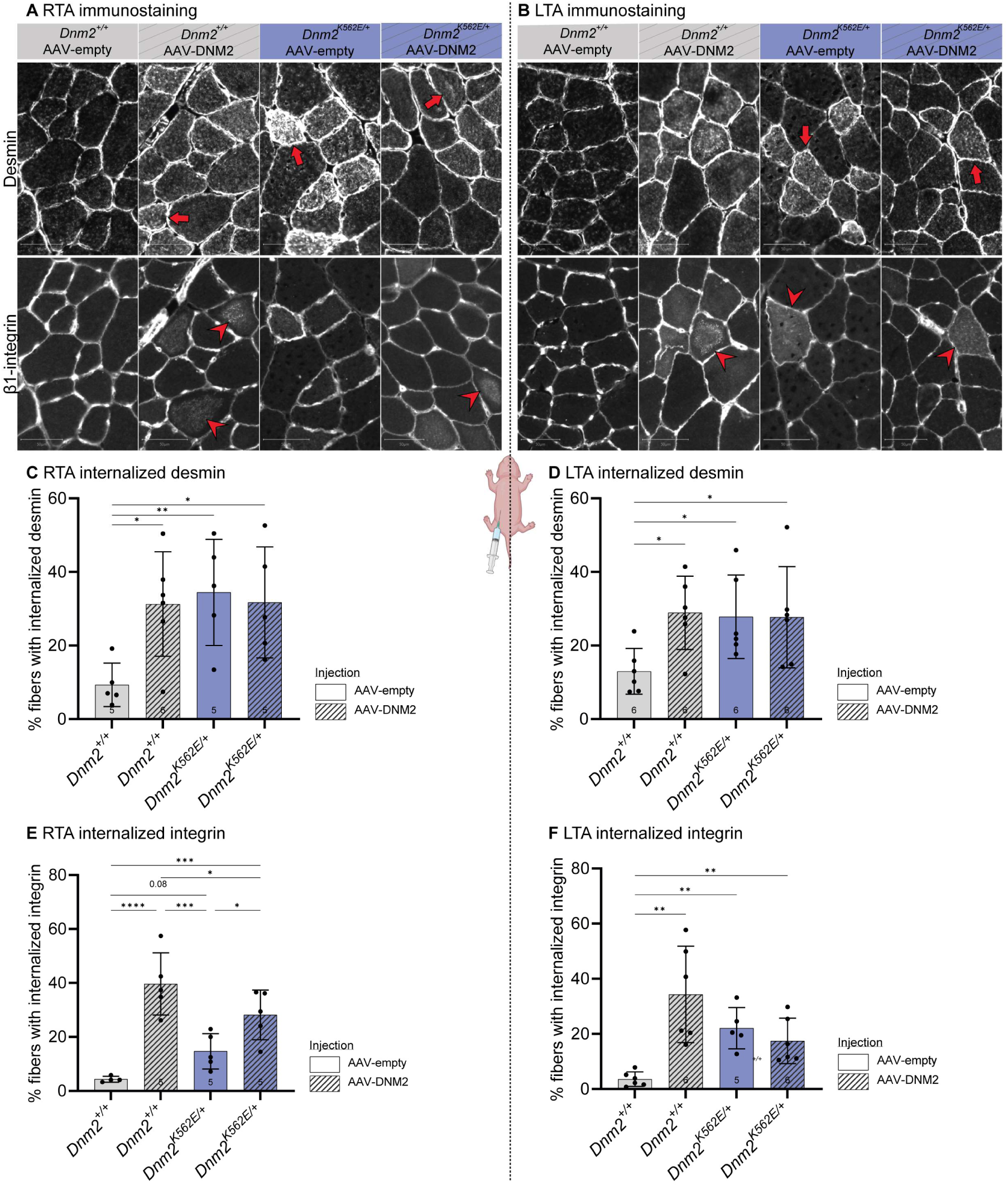
Postnatal DNM2 overexpression fails to restore desmin and integrin localization in *Dnm2-*CMT muscle. **(A-B)** Transversal sections of TA immunolabeled for desmin (top panel) or β1-integrin (second panel) in **(A)** right TA (IP-injected side) and **(B)** left TA (contralateral side). Scale bar = 50 µm. Red arrows and arrowheads mark fibers with abnormal central accumulation of desmin and β1-integrin staining respectively. **(C-D)** Proportion of fibers presenting an abnormal localization of desmin in **(C)** right TA (IP-injected side) (5≤n≤6) and **(D)** left TA (contralateral side) at 8w (n=6). **(E-F)** Proportion of fibers presenting an abnormal localization of β1-integrin in **(E)** right TA (IP-injected side) (4≤n≤5) and **(F)** left TA (contralateral side) (5≤n≤6). Each dot represents a mouse. Values are represented as mean ± SD, *p<0.05, **p<0.01, ***p<0.001, ****p<0.0001. (C-F) ANOVA test.

Overall, postnatal intraperitoneal injection of AAV-DNM2 failed to reach peripheral nerves, most likely due to the route of administration rather than the vector capsid (AAV9) or promoter (CAG), both of which have been shown to efficiently transduce nerves when delivered intravenously^33^. However, this protocol allowed to focus mainly on skeletal muscle and compare high (5.3-fold vs *Dnm2^+/+^*; 2.7-fold vs. untreated *Dnm2^K^*^562^*^E/+^*) and low (3.7-fold vs. *Dnm2^+/+^*; 2.2-fold vs. untreated) levels of DNM2 overexpression in *Dnm2^K^*^562^*^E/+^*TA muscles. Neither dose improved muscle architecture or rescued the muscle phenotypes of the *Dnm2*-CMT mouse. Instead, both levels of overexpression caused centronuclear myopathy-like abnormalities, such as organelle mislocalization, in both *Dnm2^+/+^*and *Dnm2^K^*^562^*^E/+^*mice.

## Discussion

This study investigated the contribution of skeletal muscle to the peripheral neuropathy caused by *DNM2* loss-of-function mutations and assessed the therapeutic potential of increasing DNM2 expression. We uncovered a primary involvement of skeletal muscle in *DNM2*-CMT pathology. Muscle-specific DNM2 overexpression from embryogenesis partially improved muscle organization and locomotor performance, whereas postnatal delivery failed to rescue the pathology and instead induced a CNM-like phenotype (Summary in Supplementary Fig. 5). Collectively, these findings highlight the narrow therapeutic window for safe and effective DNM2-based interventions, both in terms of developmental stage and expression level.

### A dual tissue pathomechanism in *DNM2*-related CMT

Our findings provide new insight into the dual-tissue pathomechanism of *DNM2*-related CMT. We found the *Dnm2^K^*^562^*^E/+^* mouse model exhibits muscle fiber hypotrophy, fibrosis, mislocalization of cytoskeletal and trafficking proteins, and mitochondrial dysfunction. Muscle-specific DNM2 overexpression from embryogenesis did not rescue fiber atrophy, indicating this defect may be secondary to denervation. However, it fully prevented mitochondrial defects and improved desmin and integrin expression defects, membrane trafficking, and collagen thickness, and correlated with improved locomotor activities. Notably, these features were ameliorated in the absence of nerve-targeted intervention, indicating that muscle pathology arises independently and is not merely a secondary consequence of denervation. Of note, muscle biopsies from *DNM2*-CMT patients reveal variable myopathic features. For example, Gastrocnemius biopsies (G359D mutation) show fiber atrophy, fibrosis, and type 2 fiber predominance, while Deltoid biopsies (K559del mutation) display mild oxidative changes with largely normal morphology^13,14^. In addition, the ptosis and ophthalmoplegia signs often noted in *DNM2*-CMT patients are not typical signs of CMT but are hallmarks of *DNM2*-CNM. Although chronic denervation complicates interpretation in patients, our findings strongly support a primary pathogenic role of DNM2 loss-of-function within muscle tissue itself.

### DNM2 dosage sensitivity and therapeutic window in muscle

DNM2 expression must be tightly regulated, as both loss and excess are pathogenic. Constitutive *Dnm2* knockout is embryonically lethal^28,34^, and muscle-specific deletion leads to early postnatal death^35^, highlighting the critical role of DNM2 in embryonic viability and muscle development. Conversely, DNM2 overexpression in WT animals induces centronuclear myopathy^23^, confirming dose-dependent toxicity. Elevated DNM2 levels have also been found in *DNM2-*CNM patients (unpublished data), as well as in *Dnm2-*CNM and in *Dnm2*-CMT mouse models^10^, where they correlate with muscle pathology. In our study, DNM2 overexpression in muscle (∼3.8 to 5.5-fold) following postnatal AAV injection caused CNM-like histopathology. Yet, comparable overexpression (∼4.8-fold) in *Dnm2^K^*^562^*^E/+^*;TgDNM2^SM^ improved muscular defects without toxicity. These findings collectively underscore the importance of DNM2 dosage for neuromuscular homeostasis and indicate a narrow time window for therapeutic intervention.

Our findings have important implications for therapeutic development in *DNM2*-CMT. The rationale for increasing WT DNM2 expression in a loss-of-function context is conceptually sound, to increase the overall DNM2 activity and dilute the potential dominant negative effect of the DNM2 mutant. However, our results reveal that while embryonic muscle-targeted DNM2 overexpression partially corrected muscle pathology, postnatal delivery failed to rescue disease phenotypes and, in some cases, induced CNM-like pathology. This paradox, in which efforts to compensate for a loss-of-function neuropathy risk inducing a gain-of-function myopathy, highlights that both timing and dosage are critical determinants of therapeutic efficacy and safety. The CNM-like abnormalities observed following postnatal overexpression suggest that DNM2 may have exceeded a toxic threshold in some fibers, but this is unlikely to be the major driver of pathology, as similar expression levels achieved in transgenic mice improved several phenotypes without toxicity. Thus, the lack of efficacy following postnatal injection is more likely attributable to the timing of DNM2 overexpression; postnatal expression may miss a critical developmental window or interfere with muscle postnatal maturation. Additional factors, such as the need for concurrent nerve targeting or species-specific differences in the human DNM2 isoform used here, may also contribute. It is further possible that increasing DNM2 after birth in a CMT context is simply insufficient to reverse an already established pathology. In contrast, in CNM models, DNM2 overactivity remains reversible postnatally, as antisense oligonucleotide-mediated DNM2 reduction after disease onset can rescue CNM muscle phenotypes^25^.

### Translational considerations

While muscle targeting proved effective in this model, CMT patients primarily exhibit neuropathic features. Our findings therefore raise the question of whether combined targeting of both muscle and nerve would be necessary to achieve meaningful clinical outcomes, an aspect difficult to evaluate here due to the mild nerve phenotype of the model. Such nerve targeting evaluation would require optimizing the route of AAV delivery and capsid choice for efficient nerve transduction^36,37^. Moreover, as postnatal preventive delivery failed in mice, and as patients are typically diagnosed after symptom onset, future work should assess alternative dosing strategies or timing of intervention. Given the dose-sensitive nature of DNM2, modulating its activity rather than its expression level, or employing allele-specific knockdown approaches^38^, may offer safer and more flexible therapeutic alternatives.

## Conclusions

This study demonstrates that increasing DNM2 expression specifically in skeletal muscle from embryogenesis can partially rescue key pathological and functional features of *DNM2*-CMT disease, thereby establishing muscle as a primary and therapeutically relevant target tissue in this model. Moreover, we report a first gene therapy approach aiming to overexpress DNM2 postnatally via systemic AAV delivery at birth, which instead induced CNM-like defects in muscle, underscoring the narrow therapeutic window, in terms of dose and developmental stage, for DNM2 modulation.

## Material and methods

### Mouse models

The ACTB-Cre^+^ line was on a 100% C57BL/6N background. TgDNM2^SM^ mice were generated by crossing MCK-Cre⁺ mice (expressing Cre recombinase in striated muscle from embryonic day 12.5; 100% C57BL/6N, provided by Hélène Puccio) with CAG-LSL-DNM2 mice. The latter carry a loxP-flanked STOP cassette upstream of the murine *Dnm2* gene (transcript variant 3, NM_007871.2), driven by the ubiquitous CAG promoter and inserted into the Rosa26 locus (generated at ICS; project IR8041/Kos8041; 100% C57BL/6N). Cre-mediated recombination results in muscle-specific expression of DNM2, from embryonic day 13 (E13)^39^. These mice were then crossed with *Dnm2^K^*^562^*^E/+^* knock-in mice (100% C57BL/6J; previously described^18^) to generate experimental cohorts. All mice used in the study, including controls, carried the MCK-Cre, were of mixed genetic background (50% C57BL/6N and 50% C57BL/6J), included both sexes, and were analyzed at 8 weeks of age.

For genotyping, primers “6115 Er KE” and “6116 Ef KE” were used to detect K562E mutation, primers “Cre 160” and “Cre 161” to detect the Cre, and “Sf 10692” “Wr 4035” for the CAG-LSL-*Dnm2* transgene (Supplementary Table 1 Reagents).

Mice were housed in ventilated cages, with unrestricted access to food and water. Environmental parameters were maintained at 19–22 °C temperature and 40–60% relative humidity, under a 12-hour light/dark cycle. Breeding animals were fed with SAFE® D03 diet, transitioning to SAFE® D04 after weaning.

### AAV vector design and production

Recombinant AAV9 vectors were produced by the Molecular Biology and Virus Facility at IGBMC using a standard triple transfection protocol in HEK293T/17 cells. Expression plasmid encoding human *DNM2* (transcript variant 3, NCBI RefSeq NM_004945.4) cDNA (excluding exons 12b and 13ter) was cloned under the control of the ubiquitous CAG promoter. This construct, pAAV-CAG-hDNM2 and the empty control pAAV-mU6-MCS empty, were co-transfected with the pHelper plasmid (Agilent, CA, USA) and the pAAV2/9 capsid plasmid (P0008, Penn Vector Core, PA, USA).

Viral particles were harvested 48 hours post-transfection from Benzonase-treated (100 U/mL, Merck) cell lysates. Purification was performed via iodixanol gradient ultracentrifugation (OptiPrep™, Axis Shield), followed by dialysis and concentration in Dulbecco’s PBS supplemented with 0.5 mM MgCl₂, using Amicon Ultra-15 centrifugal filters (100 kDa cutoff, Merck Millipore).

Final titers were determined by quantitative PCR (qPCR) using LightCycler 480 SYBR Green I Master mix (Roche), with primers specific to hDNM2 or the CMVe enhancer (Supplementary Table 1 Reagents). Purified vectors were aliquoted and stored at −80 °C until use.

### In vivo AAV injections

Intraperitoneal injections were performed in neonatal mice (P3-P5). A total volume of 50 µL of AAV9 vector (2.0 × 10¹² gc/mL) was injected into the lower right abdominal quadrant using a 31G syringe, corresponding to a dose of 1.0 × 10¹¹ gc per pup. For untreated controls, equivalent volumes and concentrations of AAV-empty vectors were administered using the same injection protocol.

### Behavioral tests

Behavioral assessments were conducted at 8 weeks of age, by a single experimenter blinded to genotype and treatment. The hanging test measured latency to fall from an inverted grid (max 60 s); three trials were performed per animal, and the two best were averaged. Gait analysis was performed on a motorized treadmill equipped with a camera, at a constant speed of 14 cm/s. Body stretch (nose to tail base when hind paw was in contact with the surface), stride length (step distance normalized to body stretch), and paw placement angle (angle between body axis and paw center when in contact with the surface) were quantified. Three measures were averaged for body stretch and stride, and six for paw angle (3/side). Sensorimotor coordination was assessed using the notched bar test, where mice crossed a horizontal bar with alternating notches. The number of hindlimb slips into notches were recorded over 10 trials per mouse and averaged. Body length was measured post-mortem (nose to tail base).

### Tissue collection

Mice were euthanized by carbon dioxide inhalation. TA and Soleus muscles were dissected, weighed, and snap-frozen in liquid nitrogen-cooled isopentane and stored at −80 °C for histological, RNA, or protein analyses. Sciatic nerves were snap-frozen in liquid nitrogen and stored at −80 °C.

### Muscle histology

8µm transversal TA and Soleus sections were cut using a cryostat and stained with HE, SDH, NADH, and Masson’s trichrome by the histology platform. Slides were scanned using the Carl Zeiss Axioscan 7, and 20X images were used for analysis.

Fiber segmentation was performed on HE stained sections using CellPose software^40^. MinFeret diameter were quantified in Fiji^41^. Central nuclei, SDH and NADH internalization were manually assessed using Fiji or QuPath^42^. NADH intensity was quantified as the mean gray value per fiber (scale 0–255), and the average per mouse section was plotted. Fibrotic area in Masson’s trichrome-stained sections was quantified using Fiji, adjusting color thresholds and saturation to identify blue-stained fibrosis, and normalized to the total area. One transversal section of the whole muscle was quantified for each animal.

### Muscle immunofluorescence

Transversal 8 µm TA and Soleus muscle sections were fixed in 4% paraformaldehyde for 20 minutes, permeabilized with PBS containing 0.2% Triton X-100 (Merck #T8787-250ML) for 10 minutes, and blocked for 1 hour in PBS with 0.1% Triton X-100 and 5% BSA (MP Bio #02160069-CF). Sections were incubated overnight at 4 °C with primary antibodies against integrin-β1, desmin, or collagen VI. For fiber-type identification, non-permeabilized transversal sections of Soleus muscle were incubated with antibodies against MYH7, MYH2, and MYH4 (Supplementary Table 1 Reagents). Alexa Fluor-conjugated secondary antibodies or WGA, along with DAPI, were applied for 1 hour at room temperature (RT). Slides were mounted using ProLong Gold Antifade reagent (ThermoFisher #P36934). Images were acquired at 20X magnification using either a Zeiss Axio Observer 7 microscope or the Carl Zeiss Axioscan 7. A no-primary antibody control was included for each animal.

For integrin-β1 and desmin staining in the *Dnm2^K^*^562^*^E/+^*;TgDNM2^SM^ cohort, fibers were segmented using CellPose, and mean gray intensity per fiber was quantified in Fiji. Results were represented as either intensity distribution or the percentage of high-intensity fibers. In the *Dnm2*^K562E/+^ IP-injected cohort, fiber segmentation was also performed with CellPose, and fibers were classified in QuPath as positive or negative for integrin-β1 or desmin using a fixed intensity threshold applied uniformly across samples. For WGA, fibers were segmented with Cellpose, and those with internalized staining were manually counted on Fiji. For collagen VI staining, the thickness of the signal between adjacent fibers was measured using the line tool in Fiji. Approximately 50 interfiber regions were quantified per muscle section, with six animals per genotype. Fiber types were manually quantified on Fiji and expressed as a percentage of the total number of fibers per section.

### RNA extraction and RT-qPCR

Total RNA was extracted from TA or Soleus muscles using TRI Reagent (#TR118, Molecular Research Center) and homogenized with the Precellys® Evolution Touch system (Bertin Technologies) using two 15-second pulses at 5500 rpm. Complementary DNA was synthesized from 500 ng to 1 µg of RNA using SuperScript IV Reverse Transcriptase (#18090010, Invitrogen). Quantitative PCR was carried out using SYBR Green Master Mix I (#04887352001, Roche Diagnostics) and 0.5 µM of gene-specific primers (listed in Supplementary Table 1 Reagents). Reactions were performed in technical triplicates on a LightCycler 480 system (Roche Diagnostics).

For the *Dnm2^K^*^562^*^E/+^*;TgDNM2^SM^ cohort, *Dnm2* gene expression was assessed using “*Dnm2* ex6” and “*Dnm2* ex8” primers, and normalized to the *Rpl27* housekeeping gene. In the Soleus, mtDNA was assessed by *Nd1* and nDNA by *Rps11.* For the IP-injected *Dnm2*^K562E/+^ cohort, *Dnm2* gene expression was assessed using “*Dnm2* ex10 m+h” and “*Dnm2* ex13 m+h” primers, recognizing both mouse and human sequences, and normalized to *Rps11*. Relative expression was calculated using the 2^−ΔΔCt^ method and expressed as fold change compared to the respective control group.

### Protein extraction

Muscle and nerve samples were homogenized in their respective RIPA buffers: for muscle, the buffer contained 150 mM NaCl, 50 mM Tris (pH 8), 0.5% sodium deoxycholate, 1% NP-40, and 0.1% SDS; for nerve, it contained 2% SDS, 25 mM Tris (pH 8.0), 95 mM NaCl, 2 mM EDTA, and 0.5% sodium deoxycholate. Both buffers were supplemented with 1 mM PMSF, 1 mM sodium orthovanadate, 5 mM sodium fluoride, and 1X protease inhibitor cocktail. Tissue samples were processed using a Precellys® Evolution Touch tissue homogenizer (Bertin Technologies) with two 20-second cycles at 6000 rpm and cleared by centrifugation. Protein concentrations were determined using the DC Protein Assay Kit (#5000116, BioRad).

### Western blotting

Proteins were resolved on 10% polyacrylamide gels prepared in-house according to standard protocols, except cytochrome C that was resolved on 15% gels. For muscle samples, 10 µg of protein in 12 µL were loaded per lane; for nerve samples, 7 µg of protein in 20 µL were used. Electrophoresis was performed at 130 V for approximately 1 hour. Proteins were transferred to nitrocellulose membranes using the Trans-Blot Turbo RTA Mini Nitrocellulose Transfer Kit (#170-4270, BioRad) for 5–10 minutes at 2.5 A. Membranes were stained with Ponceau S, then blocked for 1 hour in TBS containing 5% nonfat dry milk and 0.1% Tween-20 (TBST; #P2287-500ML, Merck). Membranes were incubated overnight at 4 °C with primary antibodies (Supplementary Table 1 Reagents) diluted in TBST with 5% milk. DNM2 was detected with the homemade antibody, except for the IP-injected *Dnm2*^K562E/+^ cohort (Fig. 4, Supplementary Fig. 4), where #PA5-19800 was used to detect both human and mouse DNM2. After washing, membranes were incubated for 1 hour at room temperature with the appropriate HRP-conjugated secondary antibodies. Signal detection was performed using enhanced chemiluminescence reagents (#32209, ThermoFisher Scientific), and images were acquired on an Amersham Imager 600 (GE Healthcare Life Sciences).

Band intensities were quantified using Fiji. Data were normalized to Ponceau staining and the mean value of control group, except CytC that was normalized to GAPDH instead of the Ponceau for practical reasons.

All original, uncropped western blot images used for quantification, including those not shown in the figures and their corresponding loading controls, are available in the supplementary material.

## Statistical analysis

All statistical analyses and graph generation were performed using GraphPad Prism (version 10.0.2). As no sex differences were reported in the *Dnm2*^K562E/+^ mouse model^10^, data from males and females were pooled for all analyses, except for body mass. Sex was balanced across experimental groups as much as possible (see Supplementary Table 2: Statistics). Normality was assessed using the Shapiro–Wilk test, except for large sample sizes where the D’Agostino & Pearson test was applied. For normally distributed data with equal variances, one-way ANOVA followed by uncorrected Fisher’s LSD test was used for pairwise comparisons. When variances were unequal, a Brown-Forsythe and Welch ANOVA was performed, followed by Welch’s t-tests. When data were log-normally distributed, a log transformation was applied before ANOVA analysis. For non-normally distributed data, the Kruskal–Wallis test was used, followed by uncorrected Dunn’s test. All pairwise comparisons were performed, but only statistically significant differences are shown on the graphs. A p-value < 0.05 was considered significant. Graphs display individual data points with mean ± SD. A detailed summary of the statistical tests and group sizes is provided in Supplementary Table 2.

## Ethical approval

All animal procedures were conducted in compliance with French and European regulations and were approved by the Com’Eth IGBMC-ICS institutional ethics committee (Illkirch, France) under authorization numbers APAFIS #43720-2023042415251734 and #43721-202304241436921.

## Data availability

All source data supporting the findings of this study are provided with the article.

## Acknowledgments

The authors would like to thank Ueli Suter and the Mouse Clinical Institute (ICS, Illkirch) for providing the *Dnm2^K^*^562^*^E/+^* mouse. We acknowledge the support of the scientific platforms at the Institut de Génétique et de Biologie Moléculaire et Cellulaire (IGBMC), particularly Elise Lefebvre and Paola Rossolillo for virus production, and the histology platform for performing tissue staining. We thank Théodore Bel for his help with the fiber typing. We acknowledge the IGBMC imaging center, member of the national infrastructure France-BioImaging supported by the French National Research Agency (ANR-10-INBS-04). This work of the Interdisciplinary Thematic Institute IMCBio, as part of the ITI 2021-2028 program of the University of Strasbourg, CNRS and Inserm, was supported by IdEx Unistra (ANR-10-IDEX-0002), and by SFRI-STRAT’US project (ANR-20-SFRI-0012) and EUR IMCBio (ANR-17-EURE-0023) under the framework of the French Investments for the Future Program, and by ANR CMT-GM (ANR-24-CE17-7167-01).

## Authors contribution

JL conceived and supervised the project. MG conducted all in vivo experiments. GP contributed to histological, immunofluorescence, molecular analyses and data analysis for the TgDNM2^SM^ experiments. MG conducted and analyzed all other experimental data. MG and JL wrote the manuscript.

## Bibliography

1. Laurá, M., Pipis, M., Rossor, A. M. & Reilly, M. M. Charcot–Marie–Tooth disease and related disorders: an evolving landscape. Current Opinion in Neurology 32, 641–650 (2019).

2. Praefcke, G. J. K. & McMahon, H. T. The dynamin superfamily: universal membrane tubulation and fission molecules? Nat Rev Mol Cell Biol 5, 133–147 (2004).

3. Antonny, B. et al. Membrane fission by dynamin: what we know and what we need to know. The EMBO Journal 35, 2270–2284 (2016).

4. Laiman, J., Lin, S.-S. & Liu, Y.-W. Dynamins in human diseases: differential requirement of dynamin activity in distinct tissues. Current Opinion in Cell Biology 81, 102174 (2023).

5. Züchner, S. et al. Mutations in the pleckstrin homology domain of dynamin 2 cause dominant intermediate Charcot-Marie-Tooth disease. Nat Genet 37, 289–294 (2005).

6. Durieux, A.-C., Prudhon, B., Guicheney, P. & Bitoun, M. Dynamin 2 and human diseases. J Mol Med 88, 339–350 (2010).

7. Fabrizi, G. M. et al. Two novel mutations in dynamin-2 cause axonal Charcot-Marie-Tooth disease. Neurology 69, 291–295 (2007).

8. Böhm, J. et al. Mutation spectrum in the large GTPase dynamin 2, and genotype-phenotype correlation in autosomal dominant centronuclear myopathy. Hum. Mutat. 33, 949–959 (2012).

9. Sidiropoulos, P. N. M. et al. Dynamin 2 mutations in Charcot–Marie–Tooth neuropathy highlight the importance of clathrin-mediated endocytosis in myelination. Brain 135, 1395–1411 (2012).

10. Goret, M. et al. Combining dynamin 2 myopathy and neuropathy mutations rescues both phenotypes. Nat Commun 16, (2025).

11. Fujise, K. et al. Mutant BIN1-Dynamin 2 complexes dysregulate membrane remodeling in the pathogenesis of centronuclear myopathy. Journal of Biological Chemistry 296, 100077 (2021).

12. Gallardo, E. et al. Magnetic resonance imaging findings of leg musculature in Charcot-Marie-Tooth disease type 2 due to dynamin 2 mutation. J Neurol 255, 986–992 (2008).

13. Bitoun, M. et al. A novel mutation in the dynamin 2 gene in a Charcot-Marie-Tooth type 2 patient: Clinical and pathological findings. Neuromuscular Disorders 18, 334–338 (2008).

14. Chen, S. et al. Phenotype variability and histopathological findings in patients with a novel *DNM2* mutation. Neuropathology 38, 34–40 (2018).

15. Claeys, K. G. et al. Phenotypic spectrum of dynamin 2 mutations in Charcot-Marie-Tooth neuropathy. Brain 132, 1741–1752 (2009).

16. Bolino, A. & D’Antonio, M. Recent advances in the treatment of Charcot-Marie-Tooth neuropathies. J Peripheral Nervous Sys 28, 134–149 (2023).

17. Pisciotta, C., Saveri, P. & Pareyson, D. Challenges in Treating Charcot-Marie-Tooth Disease and Related Neuropathies: Current Management and Future Perspectives. Brain Sciences 11, 1447 (2021).

18. Pereira, J. A. et al. Mice carrying an analogous heterozygous dynamin 2 K562E mutation that causes neuropathy in humans develop predominant characteristics of a primary myopathy. Human Molecular Genetics 29, 1253–1273 (2020).

19. Goret, M., Thomas, M., Edelweiss, E., Messaddeq, N. & Laporte, J. BIN1 reduction ameliorates *DNM2* -related Charcot–Marie–Tooth neuropathy. Proc. Natl. Acad. Sci. U.S.A. 122, e2419244122 (2025).

20. Laurini, C. et al. Neuromuscular disorders with evidence of both neuropathic and myopathic phenotype. Brain awaf399 (2025) doi:10.1093/brain/awaf399.

21. Bitoun, M. et al. Mutations in dynamin 2 cause dominant centronuclear myopathy. Nat Genet 37, 1207–1209 (2005).

22. Liu, N. et al. Mice lacking microRNA 133a develop dynamin 2–dependent centronuclear myopathy. J. Clin. Invest. 121, 3258–3268 (2011).

23. Cowling, B. S. et al. Increased Expression of Wild-Type or a Centronuclear Myopathy Mutant of Dynamin 2 in Skeletal Muscle of Adult Mice Leads to Structural Defects and Muscle Weakness. The American Journal of Pathology 178, 2224–2235 (2011).

24. Buono, S. et al. Reducing dynamin 2 (DNM2) rescues *DNM2* -related dominant centronuclear myopathy. Proc. Natl. Acad. Sci. U.S.A. 115, 11066–11071 (2018).

25. Muñoz, X. M. et al. Physiological impact and disease reversion for the severe form of centronuclear myopathy linked to dynamin. JCI Insight 5, e137899 (2020).

26. de Carvalho Neves, J., Moschovaki-Filippidou, F., Böhm, J. & Laporte, J. DNM2 levels normalization improves muscle phenotypes of a novel mouse model for moderate centronuclear myopathy. Molecular Therapy - Nucleic Acids 33, 321–334 (2023).

27. Moschovaki-Filippidou, F. et al. Exon skipping peptide-conjugated morpholinos downregulate dynamin 2 to rescue centronuclear myopathy. Brain awaf249 (2025) doi:10.1093/brain/awaf249.

28. Cowling, B. S. et al. Reducing dynamin 2 expression rescues X-linked centronuclear myopathy. J. Clin. Invest. 124, 1350–1363 (2014).

29. Franck, A. et al. Clathrin plaques and associated actin anchor intermediate filaments in skeletal muscle. MBoC 30, 579–590 (2019).

30. Lionello, V. M. et al. Amphiphysin 2 modulation rescues myotubular myopathy and prevents focal adhesion defects in mice. Sci. Transl. Med. 11, eaav1866 (2019).

31. Moreno-Layseca, P., Icha, J., Hamidi, H. & Ivaska, J. Integrin trafficking in cells and tissues. Nat Cell Biol 21, 122–132 (2019).

32. Bittel, D. C. et al. Secreted acid sphingomyelinase as a potential gene therapy for limb girdle muscular dystrophy 2B. Journal of Clinical Investigation 132, e141295 (2022).

33. Shen, Z. et al. Intravenous Administration of an AAV9 Vector Ubiquitously Expressing C1orf194 Gene Improved CMT-Like Neuropathy in C1orf194-/-Mice. Neurotherapeutics 20, 1835–1846 (2023).

34. Ferguson, S. et al. Coordinated Actions of Actin and BAR Proteins Upstream of Dynamin at Endocytic Clathrin-Coated Pits. Developmental Cell 17, 811–822 (2009).

35. Tinelli, E., Pereira, J. A. & Suter, U. Muscle-specific function of the centronuclear myopathy and Charcot–Marie–Tooth neuropathy-associated dynamin 2 is required for proper lipid metabolism, mitochondria, muscle fibers, neuromuscular junctions and peripheral nerves. Human Molecular Genetics 22, 4417–4429 (2013).

36. Drouyer, M. et al. Novel AAV variants with improved tropism for human Schwann cells. Molecular Therapy - Methods & Clinical Development 32, 101234 (2024).

37. Hanlon, K. S. et al. In vivo selection in non-human primates identifies AAV capsids for on-target CSF delivery to spinal cord. Molecular Therapy 32, 2584–2603 (2024).

38. Trochet, D. et al. Allele-specific silencing therapy for Dynamin 2-related dominant centronuclear myopathy. EMBO Mol Med 10, 239–253 (2018).

39. Mobley, C. B., Vechetti, I. J., Valentino, T. R. & McCarthy, J. J. CORP: Using transgenic mice to study skeletal muscle physiology. Journal of Applied Physiology 128, 1227–1239 (2020).

40. Stringer, C., Wang, T., Michaelos, M. & Pachitariu, M. Cellpose: a generalist algorithm for cellular segmentation. Nat Methods 18, 100–106 (2021).

41. Schindelin, J., et al. Fiji: an open-source platform for biological-image analysis. Nat Methods 9, 676–682 (2012).

42. Bankhead, P. et al. QuPath: Open source software for digital pathology image analysis. Sci Rep 7, 16878 (2017).

