## Supplementary material for "Muscle-specific DNM2 overexpression improves Charcot-Marie-Tooth disease in vivo and reveals a narrow therapeutic window in skeletal muscle"

**SUPPLEMENTARY MATERIALS**

Supplementary Fig. 1 to 5.

Supplementary Table 1. Reagents used. Antibodies used for immunofluorescences and western blots. Primers used for PCR and qRT-PCR.

Supplementary Table 2. Number of mice used per test and statistical analysis performed. For each test, number total of mice and sex-repartition as well as statistical analysis.

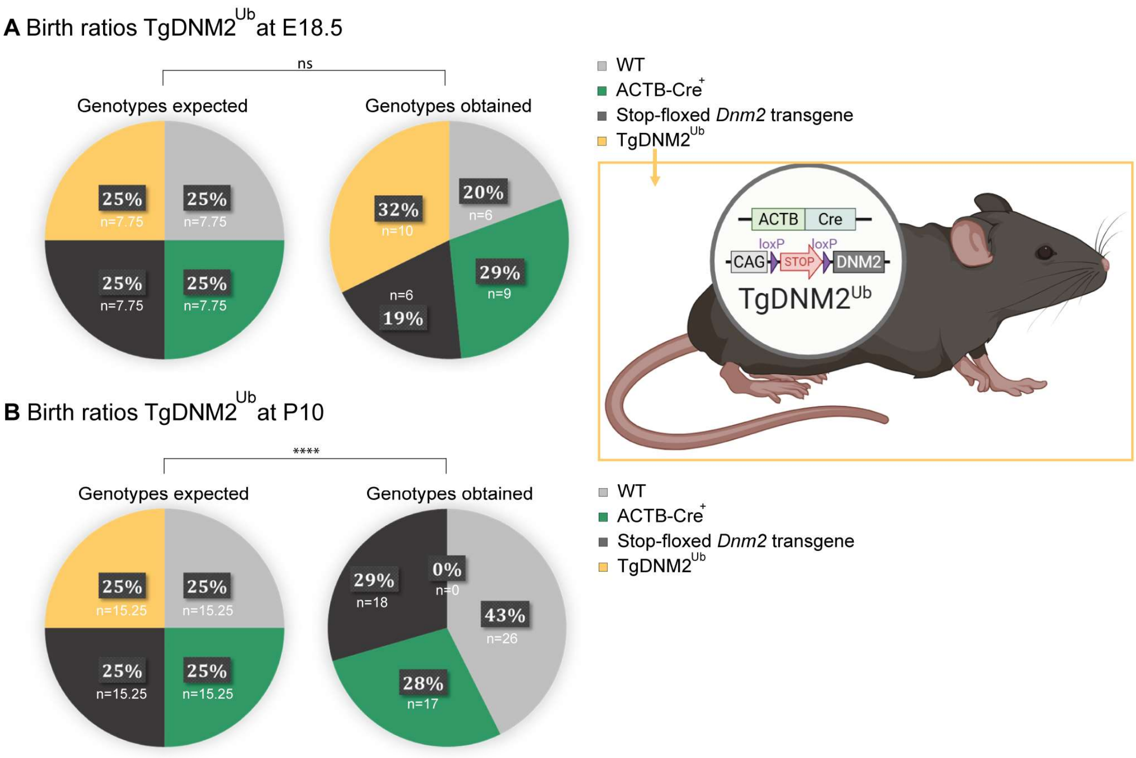


| Group Name | Description |
|---|---|
| **WT** | No Cre, no *Dnm2* transgene |
| **ACTB-Cre+** | Cre only |
| **Stop-floxed *Dnm2* transgene** | CAG-LoxP-STOP-LoxP-*Dnm2* |
| **TgDNM2$^{Ub}$** | Cre excises STOP → DNM2 overexpression |

**Supplementary Fig 1. The TgDNM2$^{Ub}$ mouse line shows perinatal lethality. (A)** Birth ratio expected and obtained at E18.5 from four different litters, in percentage and n number (n=31). **(B)** Births ratio expected and obtained at P10 for the four groups in percentage and n number (n=61). ****p<0.0001. (A-B) Chi-squared test.

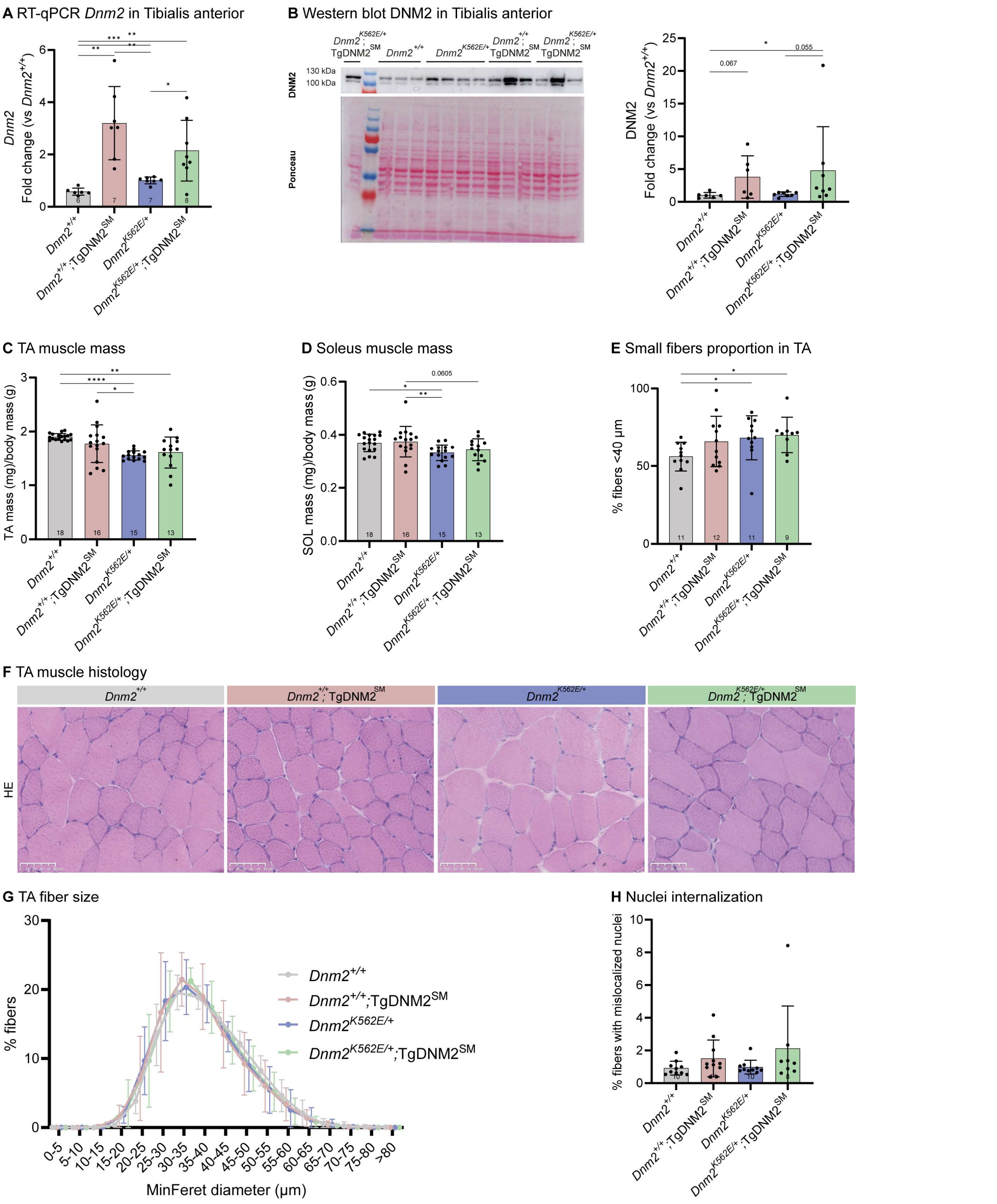


**Supplementary Fig 2. Muscle-specific DNM2 overexpression does not improve *Dnm2*-CMT muscle atrophy. (A)** RT-qPCR analysis of *Dnm2* expression in TA at 8w (6≤n≤8). **B)** Representative western blot and quantification of DNM2 protein in TA, normalized to Ponceau S staining (6≤n≤8). **(C-D)** Muscle mass of **(C)** TA, and **(D)** Soleus normalized to body mass at 8w (13≤n≤18). **(E)** Proportion of small fibers (MinFeret<40 µm) in TA sections (9≤n≤12). **(F)** TA transversal sections stained with hematoxylin-eosin (HE). Scale bar= 50 µm. **(G)** TA fibers distribution based on their MinFeret diameter (9≤n≤12). **(H)** Proportion of fibers with internalized nuclei (8≤n≤11). Each dot represents a mouse. Values are represented as mean ± SD, *p<0.05, **p<0.01, ***p<0.001, ****p<0.0001. (A-E) ANOVA test. (H) Kruskal-Wallis test.

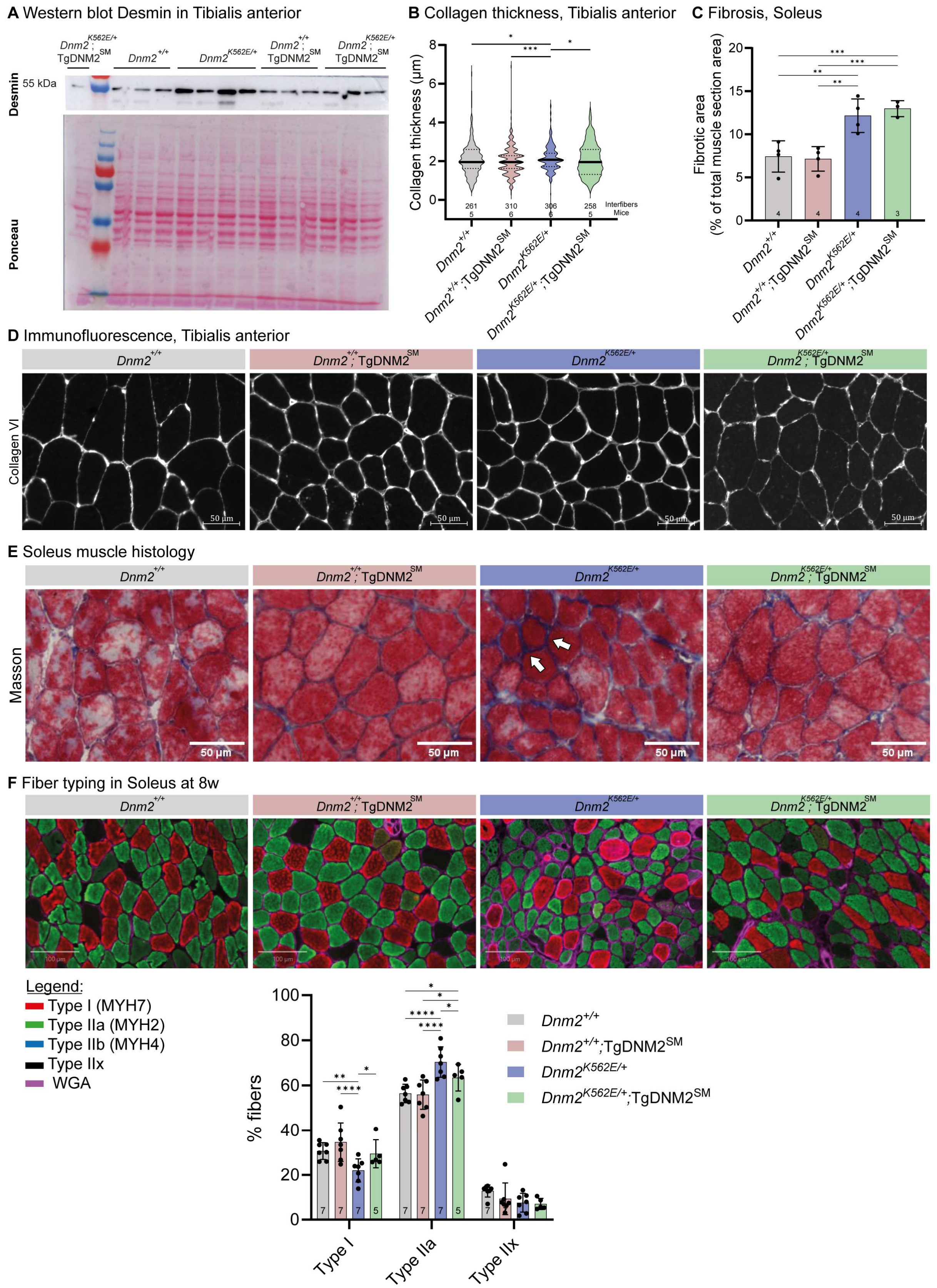


**Supplementary Fig 3. Additional data on muscle-specific DNM2 overexpression from embryogenesis: key muscle proteins expression, extracellular matrix and fiber type. (A)** Representative western blot of Desmin protein level in TA at 8w, and relative Ponceau S staining. **(B)** Collagen thickness (258≤n=interfibers≤310, 5≤n=mice≤6). **(C)** Fibrotic area in Soleus muscle (blue area on Masson's trichrome/total area) (3≤n≤4). **(D)** Immunolabeling of collagen VI in transversal TA sections. Scale bar= 50 µm. **(E)** Soleus transversal sections stained with Masson Trichrome, arrows indicate fibrosis. Scale bar= 50µm. **(F)** Immunolabeling of fiber types in transversal Soleus sections and quantification. Scale bar= 100 µm. (B) Individual interfiber spaces are plotted. (C, F) Each dot represents a mouse. Values are represented as mean ± SD, *p<0.05, **p<0.01, ***p<0.001. (B) Kruskal-Wallis test. (C, F) ANOVA test.

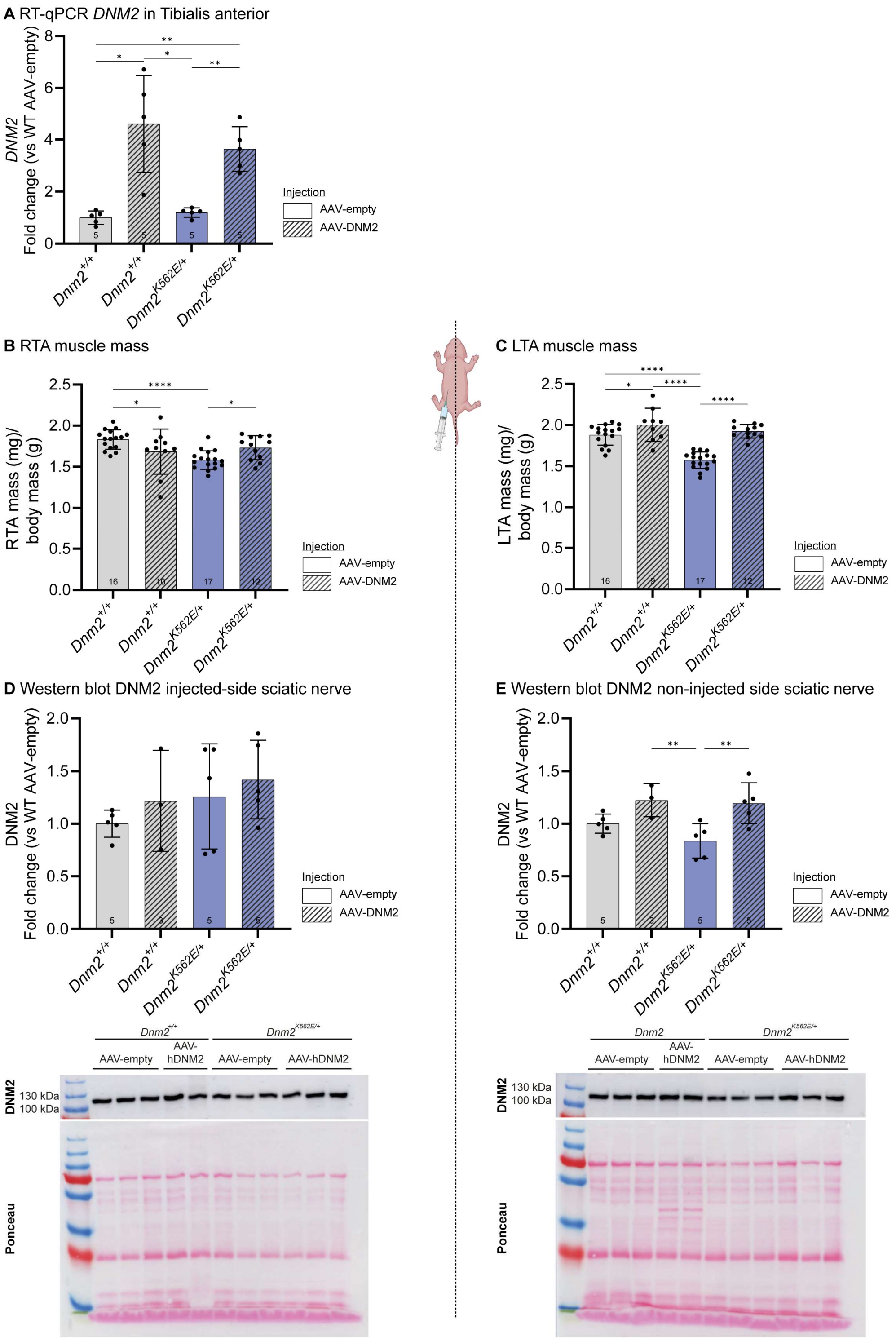


**Supplementary Fig 4. Additional data on postnatal DM2 delivery for muscle and nerves. (A)** RT-qPCR analysis of *Dnm2* expression in TA (random sides) at 8w (n=5). **(B-C)** TA muscle mass normalized to body mass at 8w: **(B)** right TA (RTA, IP-injected side) and **(C)** left TA (LTA, contralateral side) (9≤n≤17). **(D-E)** Representative western blot and quantification of DNM2 protein level in sciatic nerve at 8w, normalized to Ponceau S staining in **(D)** IP-injected side (3≤n≤5) and **(E)** contralateral side (3≤n≤5). Each dot represents a mouse. Values are represented as mean ± SD, *p<0.05, **p<0.01, ****p<0.0001. (A-E) ANOVA test.

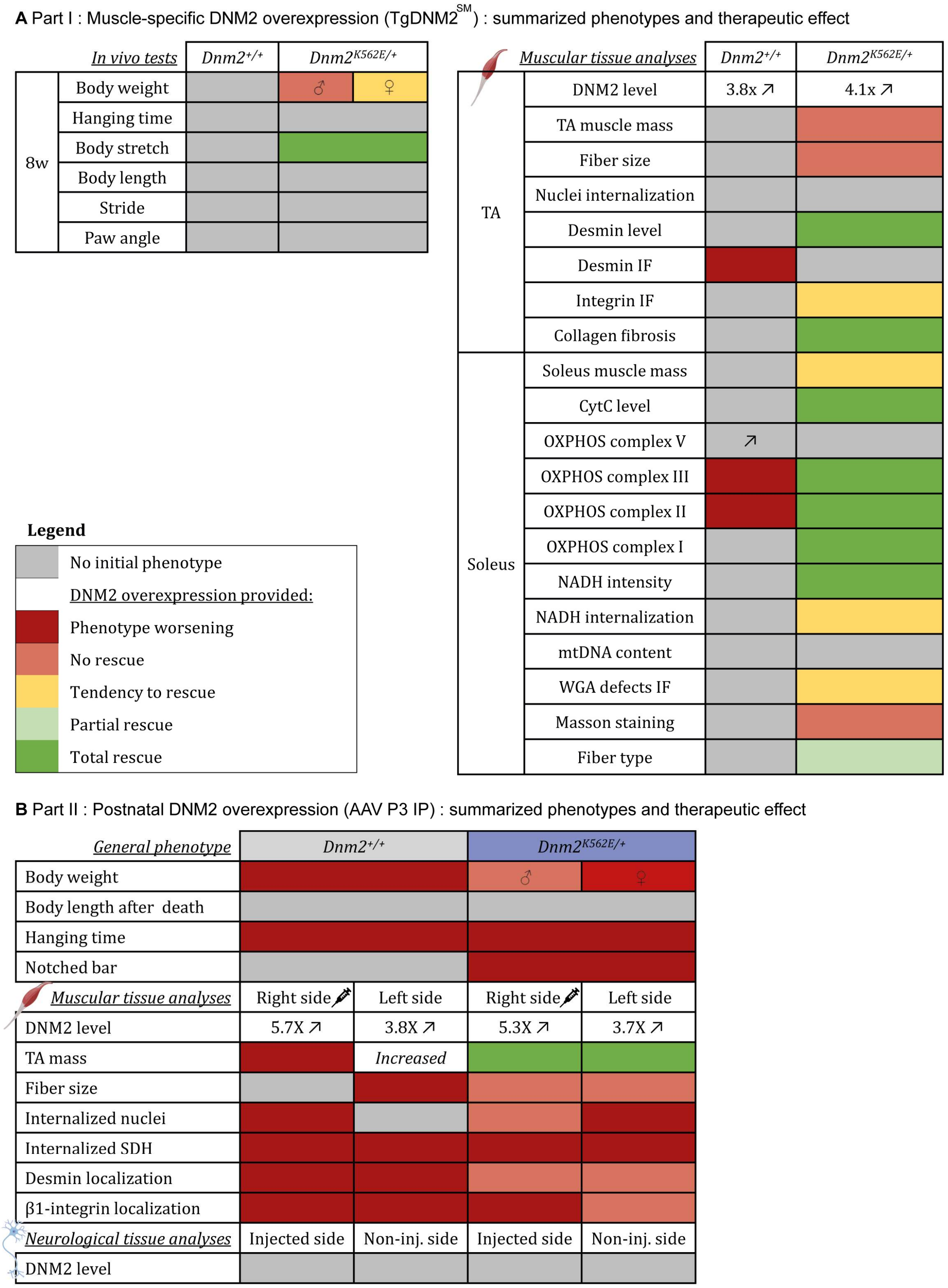


**Supplementary Fig. 5. Overview of the improvements provided by DNM2 overexpression in *Dnm2*$^{K562E/+}$ mice. (A)** Summary of the therapeutic effects from transgenic overexpression of murine DNM2 in striated muscles of *Dnm2*$^{+/+}$ and *Dnm2*$^{K562E/+}$ mice from embryogenesis, based on in vivo tests and muscle tissue analyses at 8w. **(B)** Summary outcomes following intraperitoneal (IP) injection of human DNM2 at postnatal day 3 (P3) in control and *Dnm2*$^{K562E/+}$ mice, with assessments at 8 weeks. TA= Tibialis anterior. Tendency to rescue= no significant difference between treated mutant mice and either untreated mutants or controls. Partial rescue= treated mutants differ from both untreated mutants and controls. Total rescue= treated mutants differ from untreated mutants but are similar to controls.

| Reagent/Resource | Reference or Source | Identifier or Catalog Number |
|---|---|---|
| **Muscle immunofluorescence antibodies (dilution)** | | |
| Integrin β1, rat monoclonal (1:250) | Sigma-Aldrich | MAB1997 |
| ↳ GARat Alexa 488, goat polyclonal (1:250) | Thermo Scientific | A-11006 |
| Desmin, rabbit polyclonal (1:250) | Abcam | AB15200 |
| ↳ GAR Alexa 555, goat polyclonal (1:250) | Invitrogen | A21430 |
| Collagen VI, rabbit polyclonal (1:250) | Novus Biologicals | NB 120-6588 |
| ↳ GAR Alexa 555, goat polyclonal (1:250) | Invitrogen | A21430 |
| Anti-type I fibers : MYH7 : mouse IG2b (1:50) | DSHB | BA-D5 |
| ↳ GAM IgG2b Cy3 (1 :100) | Jackson Immunoresearch | 115-165-207 |
| Anti-type IIa fibers : MYH2 : mouse IgG1 (1:50) | DSHB | SC-71 |
| ↳ GAM IgG1 Alexa 488 (1 :100) | Jackson Immunoresearch | 115-545-205 |
| Anti-type IIb fibers : MYH4 : mouse IgM (1:50) | DSHB | BF-F3 |
| ↳ GAM IgM DyLight 405 (1 :100) | Jackson Immunoresearch | 115-475-075 |
| WGA, Wheat Germ Agglutinin, Alexa 647 (1:200) | Thermo Scientific | W32466 |
| DAPI (1:1000) | / | / |
| **Western blot antibodies (dilution)** | | |
| DNM2, rabbit polyclonal (1:1000) | Homemade (2865) | N/A |
| DNM2, rabbit polyclonal (1:1000) | ThermoFisher | PA5-19800 |
| Desmin, rabbit polyclonal (1:1000) | Abcam | 15200 |
| OXPHOS, mouse monoclonal antibody cocktail (1: 1000) | ThermoFisher | 45-8199 |
| cytC, rabbit polyclonal (1:1000) | Cell Signaling | 4272 |
| GAR perox, goat polyclonal (1:10000) | Jackson Immunoresearch | 111-036-045 |
| GAM perox, goat polyclonal (1:10000) | Jackson Immunoresearch | 115-036-068 |

| **Genotyping PCR oligos** | 5’-sequence-3’ |
|---|---|
| 6115 Er KE | TACACTGTCTGCACTGTCGAGCCCTG |
| 6116 Ef KE | GCCATCTTCAACACAGAGCAGAGGTG |
| Cre 160 | GAACCTGATGGACATGTTCAGG |
| Cre 161 | AGTGCGTTCGAACGCTAGAGCCTGT |
| Sf 10692 | GGCCCACCATTATCCGCCC |
| Wr 4035 | CCTTTAAGCCTGCCCAGAAG |
| **AAV titration qPCR oligos** | |
| hDNM2 for | ATCAGGTGGACACTCTGGAGC |
| hDNM2 rev | GCATAGCTGATCTCCCGTCG |
| CMVe-CAG For5 | TACGGTAAACTGCCCACTTG |
| CMVe-CAG Rev8 | AGGAAAGTCCCATAAGGTCA |
| **qPCR oligos** | 5’-sequence-3’ |
| *Rps11* for | CGCGTGGTGAATAAGGAAGC |
| *Rps11* rev | GTAAGCACGCTCCGTCTGAA |
| *Rpl27 for* | AAGCCGTCATCGTGAAGAACA |
| *Rpl27 rev* | CTTGATCTTGGATCGCTTGGC |
| *Dnm2* ex6 | ACCCCACACTTGCAGAAAAC |
| *Dnm2* ex8 | CGCTTCTCAAAGTCCACTCC |
| *Dnm2* ex10 m+h | GTCAAGCTGAAAGAGCCCTG |
| *Dnm2* ex13 m+h | CTTCTTGTTCAGCTGCGTGC |
| *Nd1* for | AAGTTGATCGTAACGGAAGC |
| *Nd1* rev | CCCATTCGCGTTATTCTT |

**Supplementary Table 1. Reagents used.** Antibodies used for immunofluorescences and western blots. Primers used for PCR and qRT-PCR.
